# CD200 immune checkpoint reversal at the site of tumor vaccine inoculation: A novel approach to glioblastoma immunotherapy

**DOI:** 10.1101/726778

**Authors:** Zhengming Xiong, Elisabet Ampudia-Mesias, G. Elizabeth Pluhar, Susan K. Rathe, David A. Largaespada, Yuk Y. Sham, Christopher L. Moertel, Michael R. Olin

## Abstract

**Purpose:** Advances in immunotherapy have revolutionized care for some cancer patients. However, current checkpoint inhibitors are associated with significant toxicity and yield poor responses for patients with central nervous system tumors, calling into question whether cancer immunotherapy can be applied to glioblastoma multiforme. We determined that targeting the CD200 activation receptors (CD200AR) of the CD200 checkpoint with a peptide inhibitor overcomes tumor-induced immunosuppression. We have shown the clinical efficacy of the peptide inhibitor in a trial in companion dogs with spontaneous high-grade glioma; adding the peptide to autologous tumor lysate vaccines significantly increased overall survival relative to tumor lysate alone (median survival of 12.7 and 6.36 months, respectively).

**Experimental design: This study was developed to elucidate the mechanism of the CD200ARs and develop a humanized peptide inhibitor:** We developed macrophage cell lines with each of four CD200ARs knocked out to determine their binding specificity and functional responses. Using proteomics, we developed humanized peptide inhibitors to explore their effects on cytokine/chemokine response, dendritic cell maturation and CMV pp65 antigen response in CD14^+^ cells. GMP-grade peptide was further validated for activity.

**Results:** We demonstrated that the peptide specifically targets the CD200AR complex to induce an immune response. Moreover, we developed and validated our humanized peptides for inducing chemokine response, stimulating immature dendritic cell differentiation and significantly enhancing an antigen-specific response. We also determined that the use of the peptide downregulated the expression of CD200 inhibitory and PD-1 receptors.

**Conclusion:** These results support consideration of a CD200 peptide ligand as a novel platform for immunotherapy against multiple cancers including glioblastoma multiforme.

**Translational relevance:** This report evaluates the ability to modulate the CD200 immune checkpoint by employing synthetic peptides directed as ligands to its paired immune activation receptor. We previously reported the presence of CD200 in the sera and tumor vasculature of patients with glioblastoma multiforme (GBM). We have also shown that a canine CD200 activation receptor ligand extends the lives of companion dogs with high grade glioma. The data we present here show that the human peptide ligand (hCD200ARL) directed to the CD200 activation receptor on CD14+ cells activates immune upregulation through induction of a cytokine response and dendritic cell differentiation. In addition, hCD200ARL downregulates the inhibitory CD200 and PD-1 receptors. These findings provide a basis to evaluate hCD200ARL as a novel immune therapy for patients with GBM. Downregulation of PD-1 suggests that hCD200ARL may also obviate the need for PD1 and PD-L1 directed therapies for GBM and other malignancies.

## Introduction

The discovery of immune checkpoints and their inhibition (“checkpoint blockade”) is a recently developed modality for the treatment of cancer that has truly revolutionized care for some patients.^1, 2^ Current FDA-approved checkpoint inhibitors are monoclonal antibodies that can extend survival in patients with selected solid tumors such as melanoma. However, many solid tumors respond poorly to checkpoint inhibitors. This includes glioblastoma multiforme (GBM),^3^ an incurable primary central nervous system (CNS) tumor with a median overall survival of 14.6 months with the current standard of care.^4, 5^ Combinations of inhibitors that target multiple immune checkpoint pathways have been employed in an effort to significantly enhance survival. Unfortunately, these combinations may cause severe immune-related adverse events, often leading to treatment discontinuation or hospitalization, and, sometimes, death.^6,7,8,9^

The CD200 immune checkpoint results in suppression of the secretion of pro-inflammatory cytokines, including IL2 and IFNγ,^10, 11^ and increases production of myeloid derived suppressor cells (MDSCs)^12^ and T regulatory cells (Tregs),^12,13, 14^ resulting in compromised anti-tumor activity. Previously, we revealed the following mechanisms employed by the CD200 protein: (1) it is upregulated in GBM-associated endothelial cells, creating an immunological barrier around the tumor microenvironment;^10^ and (2) it is shed from tumors^12, 15^ and interacts with the inhibitory CD200 receptor (CD200R1) on immune cells in the tumor microenvironment and within the draining lymph nodes.^10, 15^ Our research focuses on the development of a therapeutic agent that targets the CD200 immune checkpoint regulatory system, which modulates the immune response through CD200R1.^10, 12^

In addition to the inhibitory CD200R1, there are a series of activation receptors (CD200AR2, 3, 4 & 5) in mice.^16^ We developed a peptide-based strategy to engage these activation receptors on immune cells^10^ and demonstrated that the inhibitory effects of CD200 protein can be surmounted by selectively engaging CD200ARs^17,11^, using specific synthetic peptide ligands (CD200AR-L) that we identified through protein sequencing and structural analyses of CD200.^10^ The ability to overpower the suppressive effects of CD200 is lost when using a scrambled CD200AR-L or CD200ARKO mice, demonstrating that these peptides mimic active sites within the CD200 protein to modulate CD200AR activity and result in immune stimulation.^10,12^

We also tested the efficacy of the CD200AR-L in companion dog spontaneous high-grade glioma using a canine-specific CD200AR-L.^18^ In this study, subcutaneous injection of the canine CD200AR-L prior to and during administration of autologous tumor lysate vaccine significantly enhanced the efficacy of the vaccine, doubling the median overall survival time compared to dogs receiving tumor lysates alone.^18^ We found the therapeutic effect of CD200-mimic peptides compelling and wished to translate these findings into the clinical setting. Herein, we describe the binding of peptide ligands to specific CD200ARs on antigen presenting cells (APCs), resulting in immune activation. Moreover, we describe the development of human CD200AR-Ls that enhance the ability of human APCs to initiate an antigen specific response.

## Materials and Methods

### Transfection

Cells from the macrophage cell line, Raw 264.7, were grown in RPMI 1640 supplemented with 10% fetal calf serum in 1% penicillin/streptomycin and incubated at 37°C until confluent. Upon confluency, transfection was performed using the Neon electroporation system (Thermo Fisher Scientific). 5 × 10^4^ cells were harvested and incubated in 10 ml Neon Buffer R with 1 ul (1ug/ul) of Clean-Cap Cas9 mRNA (TriLink Biotechnologies) and 1 ul (100 pmol/ul) CRISPR evolution sgRNA Synthego (Synthego) for 2 minutes. Following incubation, cells were placed in a Neo electroporator, 1,720 pulse voltage, 10 pulse width and 2 pulse numbers. Two days after transfection, cells were analyzed by PCR to validate the deletion.

### Immunofluorescence Cell-Binding assay

5 X10^4^ macrophage were grown in a Lab-Tek II 8 chamber slide in 200 ml RPMI containing 10% calf serum and 1% penicillin/streptomycin. Following ~70% confluency, cells were washed twice with 1X PBS and pulsed with 10uM biotinylated peptide ligand for an hour fixed in 4% PFA for 20 min at RT then incubated with streptavidin alexafluor 568 conjugate (Thermo Fisher Scientific) for 1 h, washed with 1X PBS and stained with 1ug DAPI. Imaging was performed using the Nikon Inverted TiE Deconvolution Microscope System (University of Minnesota).

### Peptide synthesis

Peptide designs were based on these regions of the CD200 protein: 1) regions of the CD200 protein previously shown to interact with CD200AR and 2) regions with the greatest homology among murine, canine, and human CD200 molecules. Three 15 amino acid peptides (P1, P3 and P4) and one 14 amino acid peptide (P2) were synthesized (Thermo Fisher Scientific, Rockford, IL). The purity of the peptides was >95% and each peptide was modified by N-terminal acetylation and C-terminal amidation to enhance their stability.

### Cytokine measurements

5×10^5^ human CD14^+^ cells were isolated from peripheral blood mononuclear cells (PBMC) using anti-CD14 beads (BD Biosciences, San Jose, CA) with a typical yield of ≥ 70% recovery and ≥ 90% purity. Cells were pulsed with 2µM each CD200AR-L (P1-4) and incubated for 48 hours. The supernatants were then analyzed by bead array for cytokine production (BD Biosciences, San Jose, CA).

### Dendritic cell differentiation

CD14^+^ cells were purified from cytomegalovirus positive (CMV^+^) HLA-A2^+^ lymphocyte packs (American Red Cross) as described above. Approximately 8×10^8^ cells were cultured in polystyrene tissue culture flasks at 37°C in 5% CO_2_. GM-CSF (25 ng/mL) and IL-4 (40 ng/mL) were added on days 3 and 5 to derive immature dendritic cells (iDCs).

### CMV assay

iDCs (5×10^5^) were pulsed with 10 µg of CMV antigen peptide pp65_495–503_ (NLVPMVATV) and cultured as described above. iDCs were then washed 3 times and co-incubated with CD8^+^ T-cells from CMV^pos^ donors (5×10^5^). CMV negative PBMCs were used as a negative control. Supernatants were collected after 48 hours of incubation and analyzed for IFN-γ production by cytometric bead array (BD Biosciences).

### Nanostring Gene Expression Analysis

Total RNA from CD14^+^ cells was sent to New Zealand Genomics Limited (Dunedin, New Zealand) to measure the expression of genes that are differentially expressed during inflammation (nCounter GX, NanoString Technologies, Seattle, WA, USA). Briefly, total RNA was extracted from CD14^+^ cells (MagJET RNA kit, Thermo Fisher Scientific, Rockford, IL) using the protocol adapted for tissue (KingFisher Duo machine, Thermo Fisher Scientific, Rockford, IL). RNA samples were then quantified (Qubit® 2.0 fluorometer, Thermo Fisher Scientific, Rockford, IL) and subjected to RNA integrity analysis (2100 Bioanalyzer, Agilent Technologies, Santa Clara, CA). Probes for the genes encoding CD44 (NM_001001392.1), NANOG (NM_024865.2), OCT4 (NM_002701.4), STAT3 (NM_139276.2), and the housekeeping genes glucuronidase beta (GUSB) (NM_000181.1), clathrin heavy chain (CLTC) (NM_4859.2), and hypoxanthine phosphoribosyltransferase 1 (NM_000194.1) were designed and manufactured by NanoString Technologies.

Expression data obtained with NanoString GX were analyzed using nSolver Analysis Software 3.0 (nanostring.com/products/nSolver) using default settings, and normalized to housekeeping genes. nSolver performed cluster analysis and generated heat maps using Java Treeview Version: 1.1.6r4. Pathway analysis was performed using PathCards Pathway Unification Database (pathcards.genecards.org)^19^. Student’s t-test was used to determine significant differences among groups (p<0.05).

## Results

### Murine CD200ARL activates CD200AR2&3 and CD200AR3&4

The CD200 checkpoint modulates immune responses through paired receptors; an inhibitory receptor (CD200R1)^20^ and activation receptors (CD200ARs). Two CD200ARs are expressed on human immune cells and four on murine cells (CD200AR2-5).^11, 21^ Although interactions between CD200 and the inhibitory receptor have been characterized, the natural ligands for the activation receptors and the molecular signaling that follows remain unknown. We have demonstrated that targeting CD200ARs may represent a promising approach for immunotherapy by enhancing an anti-glioma response in induced murine and spontaneous canine models with the addition of CD200AR-L to autologous tumor lysate vaccination. However, we wished to establish a better understanding of the mechanisms involved in targeting CD200ARs before translation to human glioblastoma patients. To achieve this, a macrophage cell line was pulsed with a fluorescently labeled murine CD200AR-L (Fig 1A). Using CRISPR, we created macrophage cell lines expressing single and duel CD200AR expressing cell lines. To achieve this, macrophage cells were first knocked out of CD200R1, or CD200AR2-4 (supplementary Figure 1a), then used to develop dual receptor knockout cell lines, then genes were knocked out to develop a single CD200R expressing cell line (supplementary figure 1b-f). Cells were validated by PCR to validate gene removal, with wildtype cells being used as a control. All receptor knockouts were sequenced to validate gene removal (supplementary figure 1a,g&h), cells with receptor 1, 3 & 4 knockouts were validated by flow cytometry, no anti-CD200AR2 is currently available. In contrast to wildtype cells (Fig. 1A), reduced peptide binding was seen on CD200AR2KO and CD200AR3KO and no peptide binding was observed on CD200AR4KO cells. Therefore, we developed cells expressing different combinations of two CD200ARs and demonstrated strong peptide binding on cells expressing CD200AR2&3, CD200AR3&4 and CD200AR2&4 (Fig. 1E&F).

**Figure 1.**
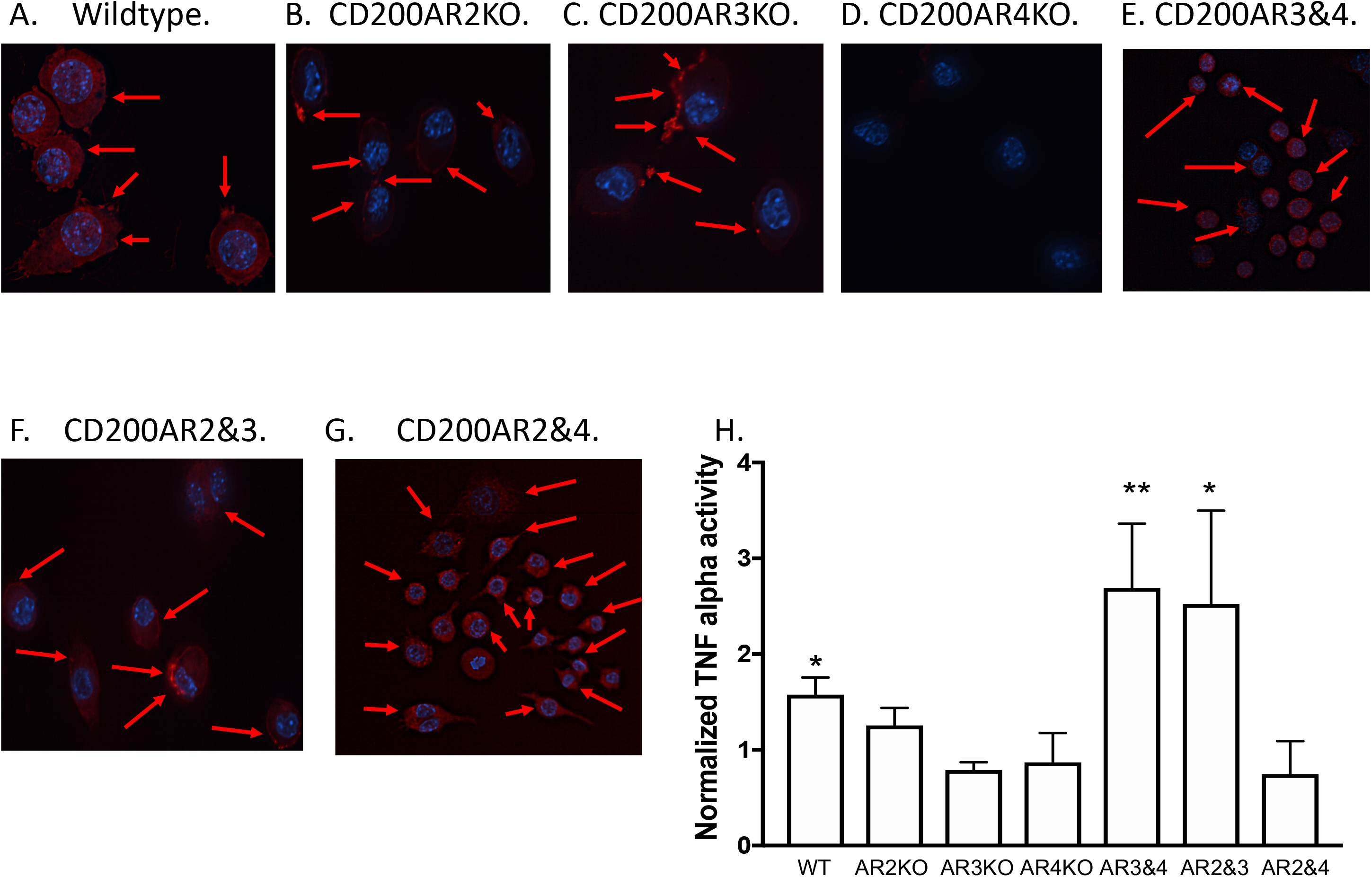
CD200AR-L binds to CD200AR complexes. (a) wildtype, (b) CD200AR2KO, (c) CD200AR3KO, (d) CD200AR4KO, (e) CD200AR3&4 expressing, (f) CD200AR2&3 expressing and (g) CD200AR2&4 expressing macrophage were pulsed with fluorescently labeled CD200AR-L and analyzed by microscopy. (H) Cells were pulsed with the non-labeled CD200AR-L and incubated for 48hrs. No pulsed cells containing the same receptors were used as a control. Supernatants were harvested and analyzed for TNFα. Error bars are representative of standard deviation (n=3/group \**P* < 0.05 and \*\**P<0.005*; by t-test

We next wanted to determine the functional effects of ligand binding to the different CD200ARs. Cell lines were pulsed with the murine CD200AR-L and supernatants were analyzed for TNFα production. These experiments correlated with the binding experiments in that only the pulsed cells expressing CD200ARs 2&&3 or 3&4 significantly produced TNFα as compared to their non-pulsed controls (Fig. G). These experiments validated that the CD200AR-L targets complexes with the CD200AR 2&3 and CD200AR3&4 to activate APCs.

### Design and identification of inhibitory peptides against CD200AR-L

Since we were now confident that we were targeting activation receptors with the peptide ligands, we sought to develop human-specific CD200AR-Ls for clinical use. Previous analysis of regions of CD200 that interact with CD200ARs revealed four regions that share significant homology among human, canine, and murine CD200 (Fig. 2A). Four CD200AR-L peptides termed P1-4 were generated. To determine whether these peptides activated human APCs as we previously observed for the murine CD200-mimic peptides,^10^ purified human CD14^+^ cells were pulsed with each of the four CD200AR-L peptides and supernatants were analyzed for immunostimulatory cytokines (Fig. 2B&C). We observed a statistically significant increase in IL-1β(p=0.0126, p=0.0364, p=0.0022, p=0.008) and TNFα(p=0.0146, p=0.0007, p=0.0002 and p=0.0082) in CD14^+^ cells pulsed with P1, P2, P3, or P4, respectively, compared to non-pulsed controls. To determine an antigen-specific response, we used a cytomegalovirus (CMV) model in which T-cells from CMV positive donors are primed with the CMV antigen pp65. Pulsing immature dendritic cells with CMV antigen pp65 + CD200AR-L peptides elicited a significant antigen-specific response, IFNγ production, compared to pp65 alone (p=0.034, p=0.033, p=0.0042 and p=0.020; P1-4 respectively, Fig. 2D).

**Figure 2.**
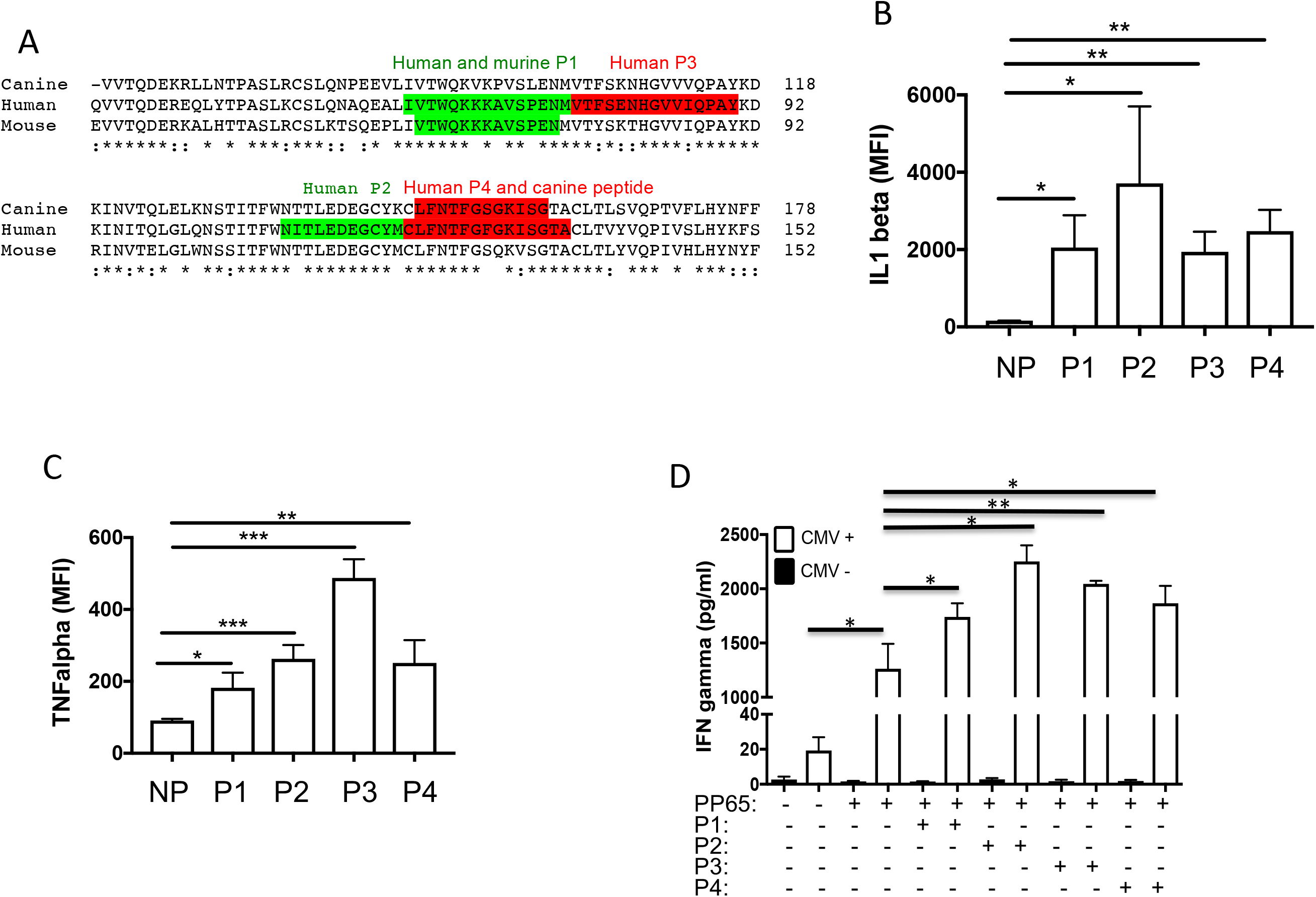
Targeting CD200 activation receptors stimulates antigen presenting cells. (a) Canine, human and murine CD200 sequences showing the homology of the different CD200 peptides. CD14^+^ cells were pulsed with peptides 1-4 and incubated for 48hrs. Non-pulsed cells were used as a control. Supernatants were harvested and analyzed for (b) IL1β and (c) TNFα. (d) Immature dendritic cells were pulsed with the CMV antigen pp65 +/− peptides 1-4. Cells were washed and autologous T cells were added back and incubated for a further 48hrs. Supernatants were harvested and analyzed for IFNγ production. Error bars are representative of standard deviation (n=3/group \**P* < 0.05, \*\**P<0.005*, \*\*\**P* < 0.0005; by t-test, \**P* < 0.05 by two way ANOVA). Data represent three separate healthy donors.

We next conducted an alanine-scanning experiment, in which each amino acid of the four peptides was substituted with alanine, with the goal of finding a peptide that induced maximal immune stimulation of APCs. Sixty-one alanine substituted peptides were created and purified CD14^+^ cells were pulsed with each peptide to determine their response, as measured by cytokine release. This led to the selection of five peptides, P1A8, P2A0, P2A5, and P4A10 (P1A8 = P1 peptide with alanine substitution of the 8^th^ amino acid), that stimulated the maximal secretion of inflammatory cytokines by CD14^+^ cells.

The effects of these peptides on a broader set of immune-stimulatory cytokines (including IL-12p70, MIG, and TNF) were then analyzed after pulsing CD14^+^ cells (Fig. 3A). In all instances, significant cytokine induction was observed after treatment with the five peptides. To further characterize the effect of these peptides on CD14^+^ cells, NanoString analysis was performed. Consistent with our previous observation that CD200 exposure suppresses TNF signaling in APCs and CD200-mimic peptides reverse this effect, P1A8, P2A0, and P4A10 treatment induced a notable increase in the mRNA expression of cytokines released in response to the TNF signaling pathway (Table 1, Fig. 3A). These results were recapitulated using a NanoString platform designed to detect the mRNA expression of TNF regulated cytokines (Fig. 3B). The three most potent peptides that consistently induced the mRNA expression of TNF regulated cytokines (P1A8, P2A0, and P4A10) were selected for subsequent analysis.

**Figure 3.**
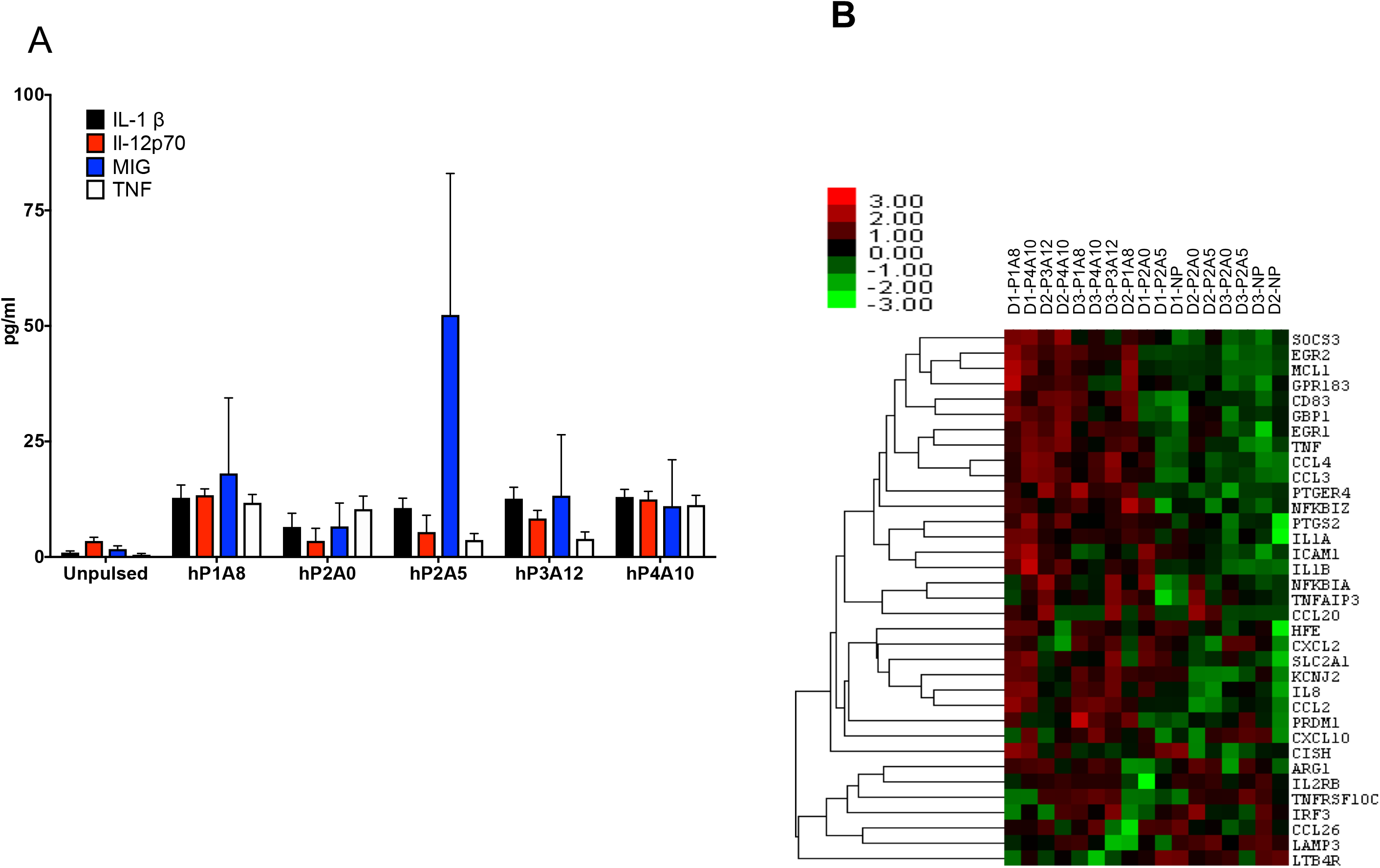
Alanine substitutions enhance antigen-presenting cell stimulation. CD14^+^ cells were pulsed with equal molar ratio of peptides P1A8, P2A0, P2A5, P3A12 and P4A10, and (a) incubated for 48hrs, Supernatants were harvested and analyzed for IFN-1β IL-12p70, MIG and TNFα production. (b) In separate experiments, cells were incubated for 1hr. RNA was harvested and analyzed by NanoSight for immune related transcription alterations. Results were analyzed by nSolver. Pulsed cells were normalized to a non-pulsed (NP) control to derive heat map. Cluster analysis of the 35 genes show significant expression changes in one or more of the treated samples when compared to the NP controls. Excludes the outlier D1-P3A12. Error bars are representative of standard deviation (n=3 donors run in triplicate \**P* < 0.05, \*\**P<0.005*, \*\*\**P* < 0.0005; by t-test, \**P* < 0.05 by two way ANOVA). Heatmap was generated by nSolver using Java Treeview.

**Table 1.**
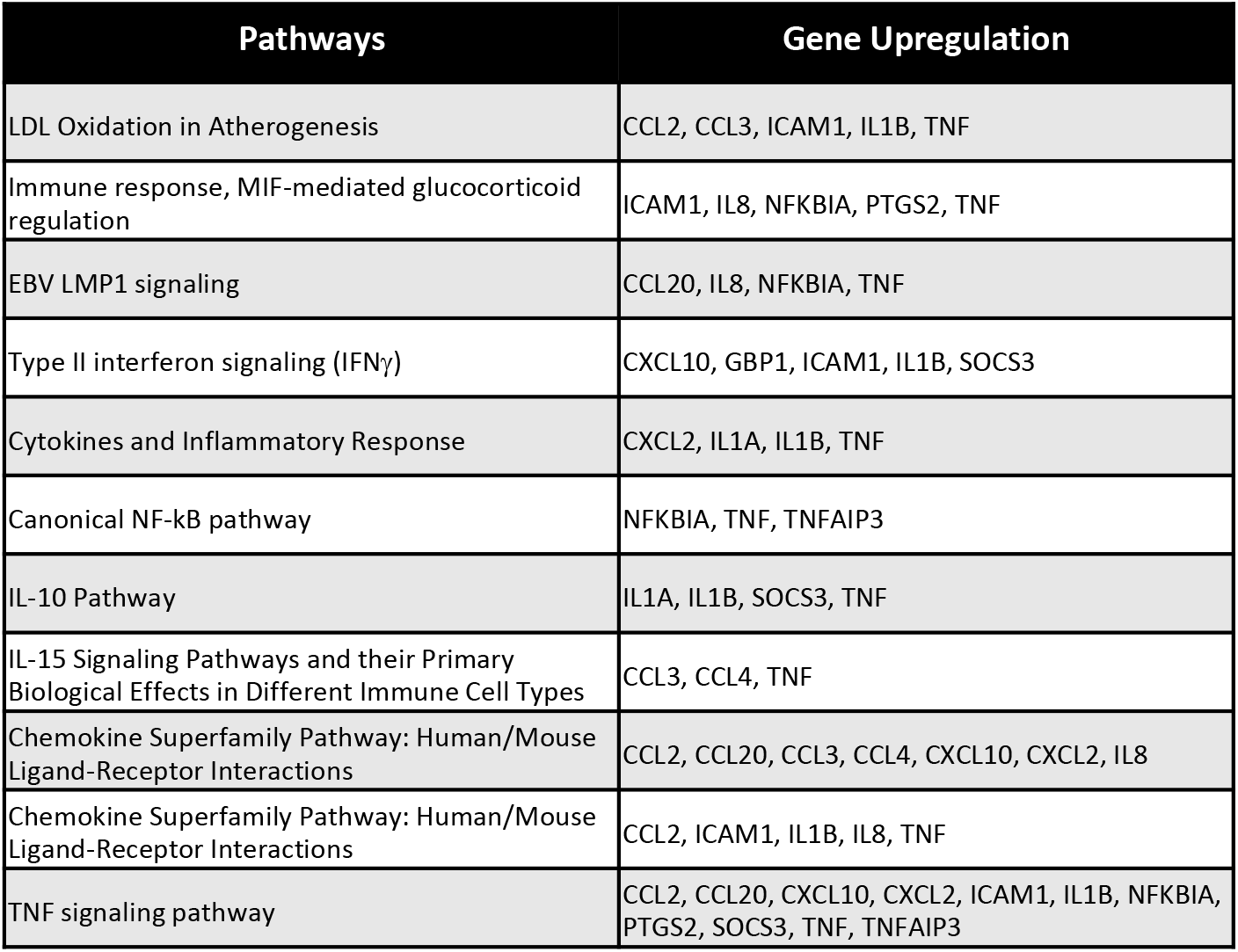
Pulsed and non-pulsed CD14 cells were analyzed by IPA analysis for upregulation of the TNF pathway.

| Pathways | Gene Upregulation |
| --- | --- |
| LDL Oxidation in Atherogenesis | CCL2, CCL3, ICAM1, IL1B, TNF |
| Immune response, MIF-mediated glucocorticoid regulation | ICAM1, IL8, NFKBIA, PTGS2, TNF |
| EBV LMP1 signaling | CCL20, IL8, NFKBIA, TNF |
| Type II interferon signaling (IFNγ) | CXCL10, GBP1, ICAM1, IL1B, SOCS3 |
| Cytokines and Inflammatory Response | CXCL2, IL1A, IL1B, TNF |
| Canonical NF-kB pathway | NFKBIA, TNF, TNFAIP3 |
| IL-10 Pathway | IL1A, IL1B, SOCS3, TNF |
| IL-15 Signaling Pathways and their Primary Biological Effects in Different Immune Cell Types | CCL3, CCL4, TNF |
| Chemokine Superfamily Pathway: Human/Mouse Ligand-Receptor Interactions | CCL2, CCL20, CCL3, CCL4, CXCL10, CXCL2, IL8 |
| Chemokine Superfamily Pathway: Human/Mouse Ligand-Receptor Interactions | CCL2, ICAM1, IL1B, IL8, TNF |
| TNF signaling pathway | CCL2, CCL20, CXCL10, CXCL2, ICAM1, IL1B, NFKBIA, PTGS2, SOCS3, TNF, TNFAIP3 |

### Targeting CD200 activation receptors enhances dendritic cell maturation

**The** NanoString analysis described above suggests that P1A8, P2A0, and P4A10 induce expression of genes implicated in dendritic cell maturation (Table 1). To substantiate this observation, CD14^+^ cells were isolated from healthy human donors and pulsed either with GM-CSF + IL4 or P1A8, P2A0 or P4A10. These studies demonstrate that the CD200AR-Ls induce the differentiation of CD14 cells to immature dendritic (iDC) cell populations with decreasing CD14 expression and increased expression of co-stimulatory molecules CD80/86 and HLA-DR, compared to cells treated with GM-CSF + IL4 alone (p< 0.0001) (Fig. 4A). Moreover, we observed synergistic upregulation of CD80/86 and HLA-DR when CD14+ cells were incubated with GM-CSF + IL4 and each of the peptides (p<0.0001) (Fig. 4B). These results suggest that P1A8, P2A0, and P4A10 enhance differentiation of CD14^+^ monocytes from healthy donors into iDCs for antigen priming.

**Figure 4.**
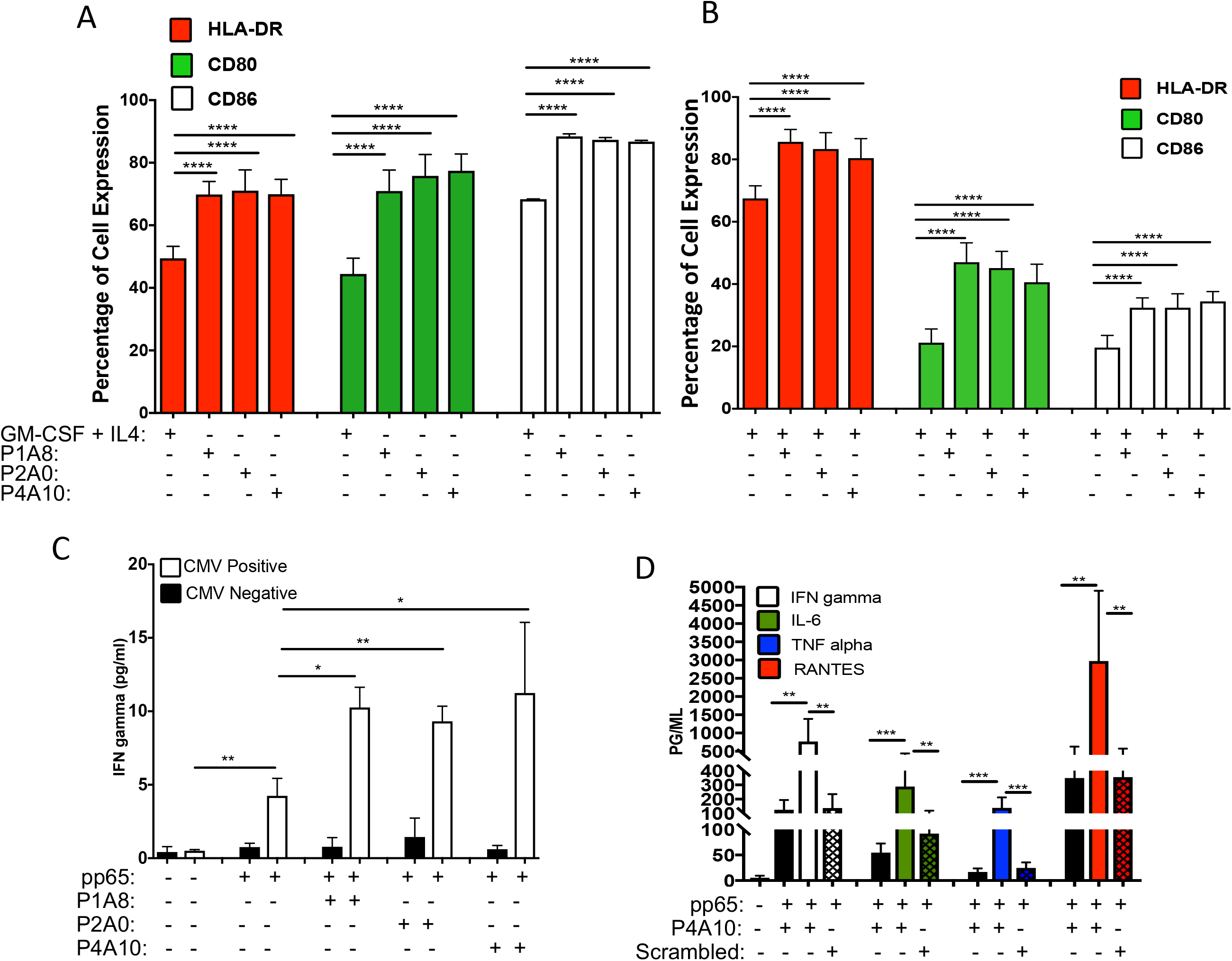
Targeting CD200 activation receptors enhances dendritic cell maturation. CD14 purified cells were pulsed with (a) GM-CSF + IL4 or equal molar ratios of peptides P1A8, P2A0 or P4A10 or (b) GM-CSF + IL4 with equal molar ratios of peptides P1A8, P2A0 or P4A10 and incubated for 48hrs. Cells were harvested and phenotyped for CD80, CD86 and HLA-DR. (c) Immature dendritic cells were pulsed with the CMV antigen pp65 +/− equal molar ratios of peptides P1A8, P2A0 or P4A10. Cells were washed and autologous T-cells were added back and incubated for a further 48hrs. Supernatants were analyzed for IFNγproduction. (d) GM-CSF + IL-4 derived immature dendritic cells were pulsed the CMV antigen pp65 +/− P4A10. Cells were washed, autologous T-cells were added back and incubated for a further 48hrs. Supernatants were analyzed for IFNγ IL-6, TNFα and RANTES production. Error bars are representative of standard deviation (n=3 donors each run in triplicate \**P* < 0.05, \*\**P<0.005*, \*\*\**P* < 0.0005; by t-test).

To assess the effects of the peptides on an antigen-specific response, GM-CSF + IL4 induced iDCs were pulsed with the CMV antigen pp65 with and without an equimolar ratio of P1A8, P2A0, or P4A10. Autologous T-cells were incubated with the DCs for 48 hours and IFNγ production was then measured in supernatants. pp65 exposure increased T-cell IFNγ production approximately four-fold (Fig. 4C). Co-treatment of immature dendritic cells with P1A8, P2A0, or P4A10 and pp65 induced another 2-3 fold increase in IFNγrelease, demonstrating that P1A8, P2A0, and P4A10 enhanced the dendritic cells’ ability to induce human T-cell antigen specific response. We next tested the effects of P4A10, the most potent of the three peptides, on T-cell secretion of inflammatory cytokines after pp65 antigen presentation by dendritic cells. We observed that P4A10 peptide treatment of dendritic cells enhanced T-cell secretion of IFNγ, IL6, TNFα, and RANTES by 5.4, 5.6, 16.6 and 16.3 fold, respectively. A scrambled peptide control failed to enhance the pp65 response (Fig. 4D). These results show that P4A10 enhanced the ability of APCs to induce a T-cell mediated immune response.

### Validation of the GMP peptide

Due to formulation stability issues of P4A10, we decided to take the human P1A8 peptide, which showed efficacy in our murine survival model, forward to a phase I clinical trial. To insure the GMP grade peptide retained activity following development and subsequent vialing formulation, we compared its binding kinetics to that of the murine peptide on HEK293 cells (Fig. 5A). Next, we tested the peptide using a human CMV model to assess an anti-pp65 response, demonstrating that the GMP CD200AR-L maintained its ability to enhance an antigen specific response (Fig. 5B). Finally, we observed the binding kinetics of the GMP-grade peptide using CD14 and found a dose response peaking around 1500uM (Fig. 5C&D). All results confirm maintenance of immunostimulatory activity of the GMP grade peptide.

**Figure 5.**
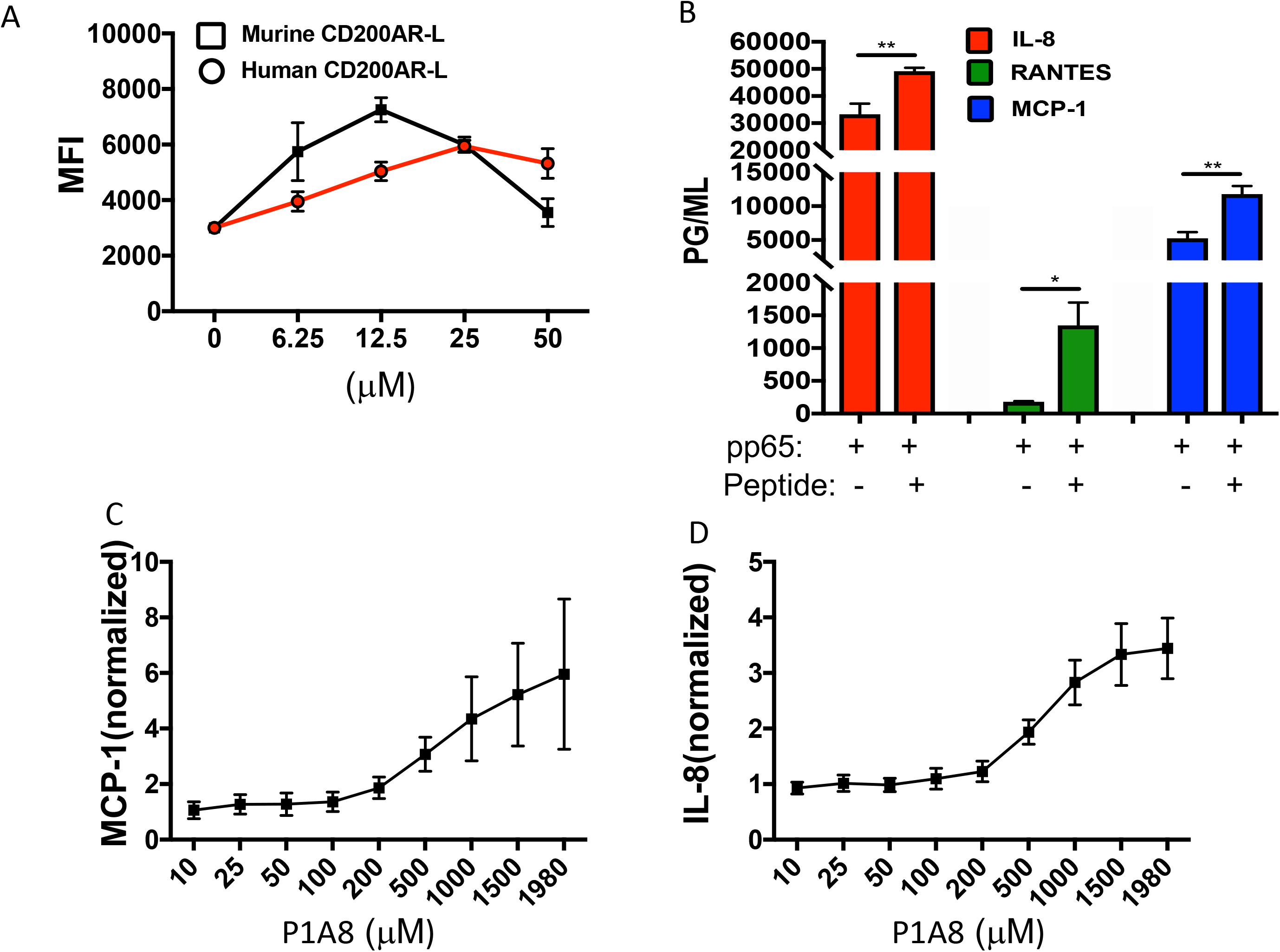
Validation of the GMP peptide. (a) HEK293 cells were pulsed with the murine P1A12 and human P1A8 fluorescently labeled CD200AR-L and analyzed for binding. (b) GM-CSF + IL-4 derived immature dendritic cells were pulsed the CMV antigen pp65 +/− P1A8. Cells were washed, autologous T-cells were added back and incubated for a further 48hrs. Supernatants were analyzed for IFNγ production. CD14+ cells were pulsed with different concentrations of P1A8 and analyzed for (c) MCP-1 and (d) IL-8 production. Cells were normalized to non-pulsed cells. (n=3 donors run in triplicate \**P* < 0.05, \*\**P<0.005*, \*\*\**P* < 0.0005; by t-test).

### Antigen presenting cells primed with P1A8 down-regulate expression of the inhibitory CD200R1

Targeting the CD200ARs activates the immune system by overpowering the suppressive effects of CD200. To gain a better understanding of this mechanism, we pulsed the human macrophage THP-1 and CD14 cells with the GMP grade P1A8. We showed that this treatment decreased the expression of the inhibitory receptor (Figs. 6A-C), which should render these cells resistant to the effects of CD200 solubilized to the draining lymph nodes or the glioblastoma microenvironment. This suppression was not observed in mock-treated control reactions. Interestingly, downregulation of PD-1 on APCs was also observed (p=0.005) (Fig. 6D&E). These results open the possibility of overcoming CNS immunosuppression through modulation occurring outside of the CNS.

**Figure 6.**
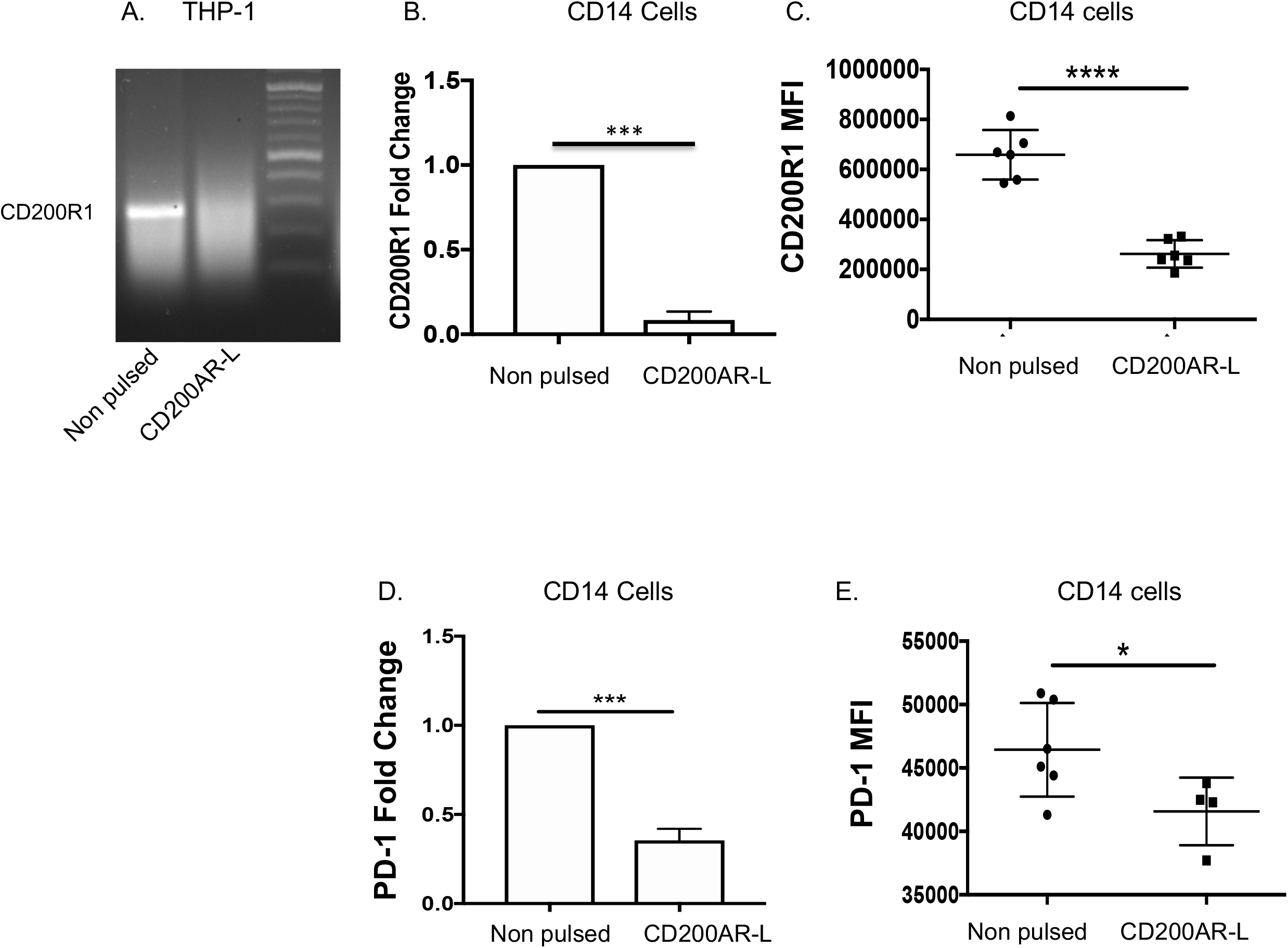
Inhibitory CD200R1 is downregulated following pulsing with the CD200AR-L. (a) Human monocyte cell line THP-1 cells were pulsed with the human CD200AR-L (P1A8), following 24hr incubation, cells were analyzed by western analysis for inhibitory CD200R1 expression. CD14^+^ cells were pulsed with the CD200AR-L and analyzed for (b) CD200AR-L transcriptional changes and (c) protein levels by flow cytometry. CD14^+^ cells were pulsed with the CD200AR-L and analyzed for (d) PD-1 transcriptional changes and (e) protein levels by flow cytometry. (n=3 donors run in triplicate \**P* < 0.05, \*\**P<0.005*, \*\*\**P* < 0.0005; by t-test).

## Discussion

Immune checkpoint inhibitors are currently at the forefront of developing immunotherapies.^22, 23^ The most clinically successful have been those against cytotoxic-T-lymphocyte-associated protein 4 (CTLA4) and the paired programmed cell death 1 receptor (PD-1)/programmed cell death ligand 1 (PD-L1). Nevertheless, tumors can inhibit the anti-tumor immune response through multiple checkpoints, hindering the use of these inhibitors as monotherapy. This is particularly critical for high-grade malignant brain tumors with lower mutational burden and/or lower immunogenicity. Therefore, combination therapies that may significantly enhance survival are often used concomitantly. However, this frequently causes serious immune-related adverse events.^6,7^

No single currently approved checkpoint inhibitor has demonstrated significant efficacy in high-grade glioma patients. Here, we explored an alternate paradigm with immune checkpoint reversal at the site of autologous tumor vaccine inoculation outside of the CNS. Our previous studies provided compelling evidence that the CD200 immune checkpoint protein in tumor lysates suppresses the ability of APCs to trigger an effective anti-tumor immune response through locally recruited T-cells.^10, 12^

Several rigorous studies have provided evidence that targeting the CD200 checkpoint enhances immunotherapy.^24–28^ In the most advanced of these studies, a monoclonal antibody against CD200 (ALXN6000) was evaluated in a clinical trial initiated in 2008 for relapsed or refractory B-cell chronic lymphocytic leukemia (B-CLL) and multiple myeloma (NCT00648739) patients. The results for B-CLL were posted on the Alexion website, but those for multiple myeloma were not.

We believe that there are problems associated with the use of an anti-CD200 antibody for GBM, some of which are exemplified by the Alexion results: i) antibodies fail to cross the blood-brain barrier, which limits their efficacy in CNS tumors; ii) multiple cells including neurons and immune cells express CD200^29, 30^, therefore, the use of an anti-CD200 antibody might cause off-target toxicity and decrease effectiveness. Alexion reported that in B-CLL patients, 95% of the patients had up to 98% reduction of CD200^+^CD4^+^ T cells. The loss of these immune cells may create an immunocompromised condition and may be the reason that only 36% of the patients had a 10% reduction in bulk disease. Although they reported CD200^+^ B-CLL was reduced 64%-75% in 67% of patients, CD200 is secreted from tumors^31^ and this parameter does not correlate with tumor reduction.

We chose to develop peptide ligands to target the CD200 activating receptors on APCs. Peptides have the ability to penetrate further into tissue^32^ and have higher activity per unit mass, greater stability and reduced potential for nonspecific binding that may result in decreased toxicity.^33^ Having clearly demonstrated efficacy with the use of a synthetic peptide ligand, the mechanism of immune response modulation through binding to the activation receptors was still unknown. Therefore, we developed three murine CD200AR-Ls that demonstrated different survival results. The specific murine ligand, P1, which was predicted to bind primarily to CD200AR4,^12^ enhanced survival in our glioma model, whereas the ligands predicted to bind primarily to CD200AR2 and CD200AR3 enhanced survival in our breast tumor model but failed in our glioma model.^12^ In these studies, we are interested in the role of activation receptors (CD200ARs) therefore, we focused only on those cell lines expressing CD200AR2-4. We observed that cells expressing CD200AR2&3 and 3&4 responded to peptide stimulation by the P1 ligand, while cells expressing receptors 1,2&3, 1,3&4 or 2,3&4 failed to bind the florescent P1 or produce TNF alpha. In addition, P1 ligand binds to the CD200AR2&4 cell line, however, no TNF production was observed. We suggest the these receptors are producing different immune response, a phenomenon that is currently under investigation in our laboratory. These studies lead to the hypothesis that the activation receptors (CD200ARs) function by creating complexes to modulate immune activation. This would explain our observation that targeting different CD200ARs induced different survival benefits in our breast carcinoma and glioma murine models.^12^

Our studies provide compelling data that the presence of the CD200 protein in a brain tumor lysate suppresses the capacity of local APCs to activate recruited T-cells and trigger an effective anti-tumor immune response.^10, 12^ We have built on this observation and tested peptides that target CD200 mediated immunosuppression, successfully reversing the immunosuppressive effect of CD200 in murine studies.

While murine brain tumor models have yielded valuable insights into the etiology of glioblastoma, novel therapies that showed enormous promise in these models have frequently failed in clinical studies. Recently, attention has been focused on companion dogs as a translational model due to their strong anatomical and physiological similarities to humans and the sheer number of pet dogs that are diagnosed and managed with cancer each year.^34–36^ Strong similarities have been shown between the canine and human genome, especially with respect to gene families associated with cancer. These combined factors suggest cancer in companion dogs as a viable model for pre-clinical human cancer research including brain tumors.^37–39^ Due to the success in the canine CD200 trial,^40^ the human CD200AR-L, P4A10, analogous to the canine CD200AR-L, was selected for use in a human phase I trial. However, the charges within this peptide made it difficult to scale up for GMP production. We therefore chose P1A8 based on results from our murine studies. Our final human *in vitro* studies demonstrated that hP1A8 enhanced DC maturation and cytokine production as well as an antigen-specific response. In addition to cell activation, we also demonstrated downregulation of the inhibitory receptor, CD200R1. This is important because CD200 secreted by the tumor suppresses the ability to mount an anti-tumor response and CD200 is upregulated in the tumor-associated vascular endothelium (as evidenced in our earlier studies)^10^, limiting the ability of immune cells to extravasate in response to immunotherapy due to the binding of CD200/CD200R1 on immune cells. We believe downregulation of CD200R1 allows immune cells to move into the tumor microenvironment from the tumor vasculature. Moreover, the downregulation of inhibitory CD200R1 and PD-L1 renders the immune cells resistant to tumor-induced suppression within the tumor microenvironment. We hypothesize that the significant survival response seen in the canine pre-clinical trial is due to the ability of the CD200AR-L peptide to override the suppressive effects of multiple immune checkpoints.

---

**Supplementary figure 1.**
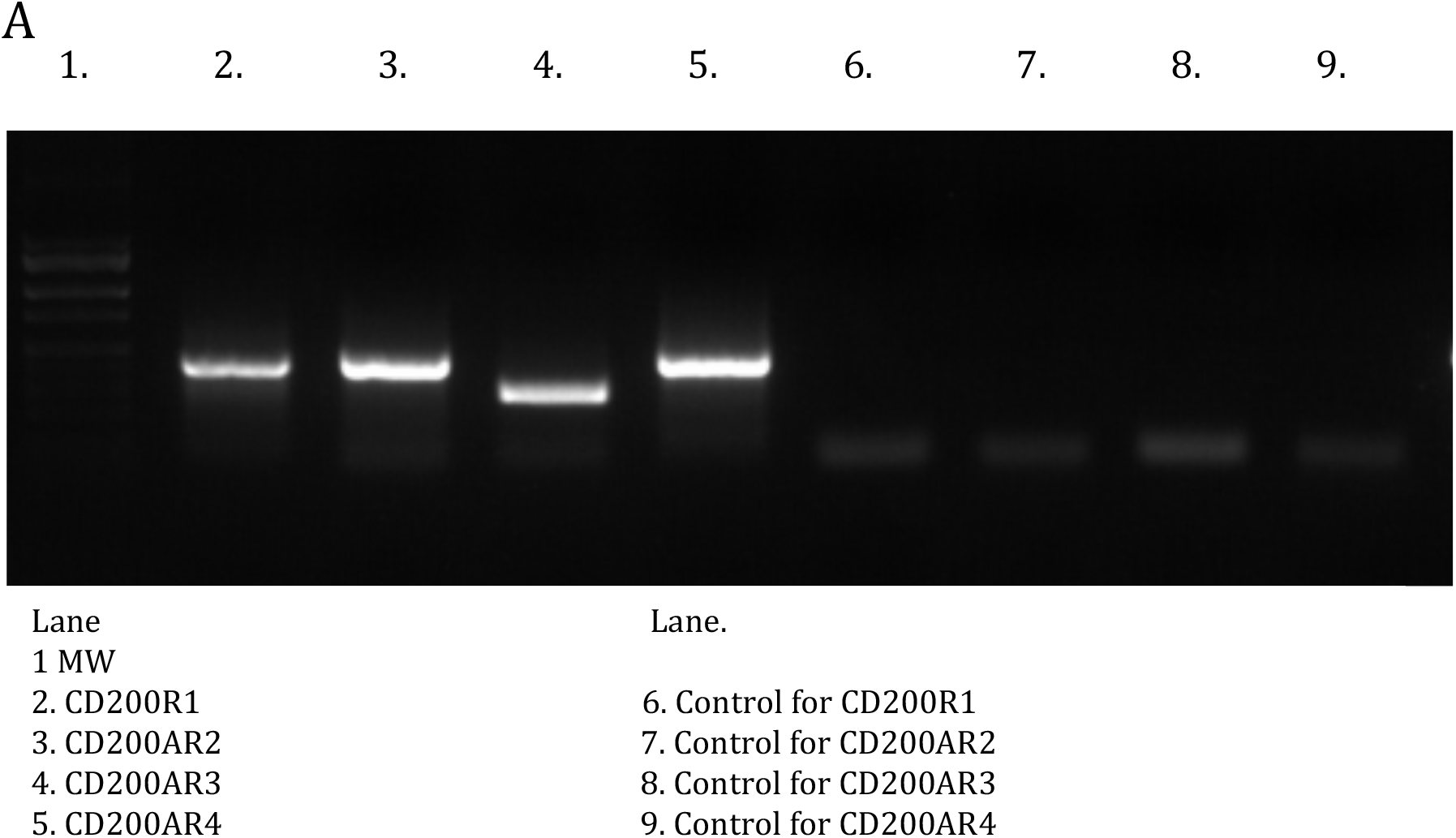
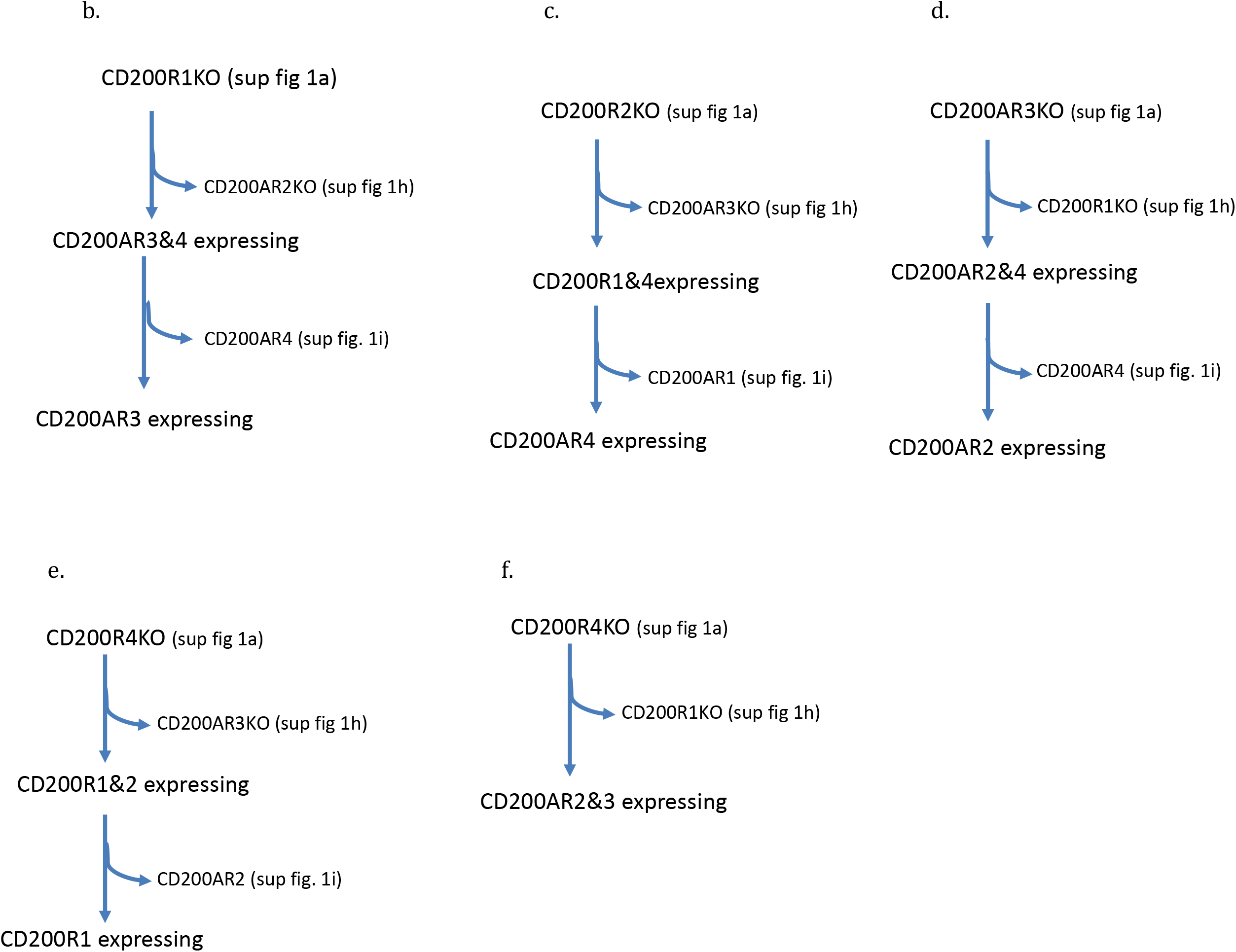
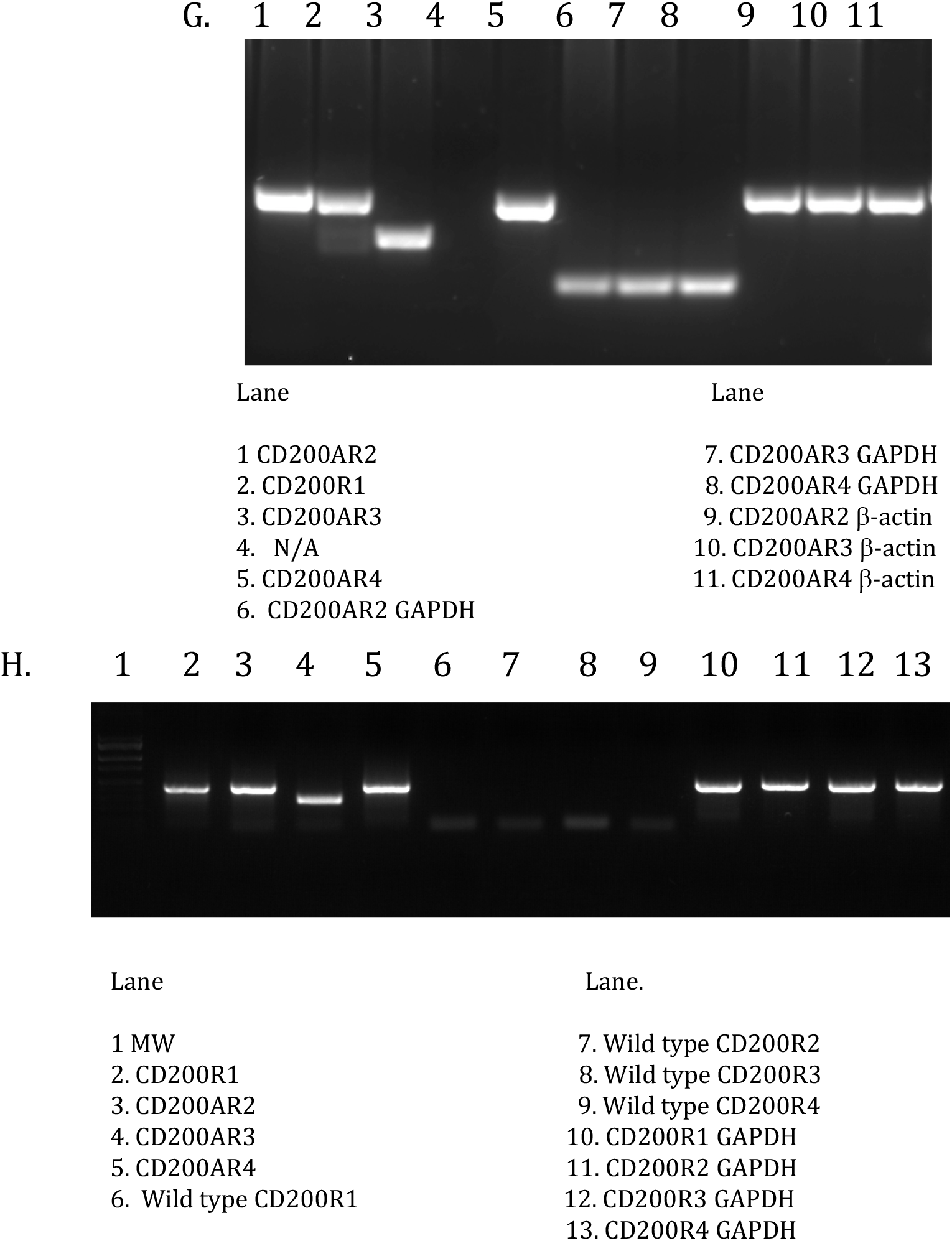
Generation of Knockout Macrophages (Raw 264.7). (a) Monocyte raw 264.7 cell line was used to knockout single CD200 receptor (CD2001KO, CD200AR2KO, CD200AR3KO or CD200AR4). (b) CD200R1 knockout line was used to derive CD200R1&2 KO (CD200AR3&4 expressing) cells, then further used to derive CD200AR3 single expressing cell line. (c) CD200R2KO cells were used to derive CD200AR2&3KO (CD200R1&4 expressing) cells, then used to derive CD200AR4 single expressing cells. (d) CD200AR3KO cells were used to derived CD200R1&3KO (CD200AR2&4 expressing) cells, then further used to derive CD200AR2 single expressing cells. (e) CD200R4KO cells were used to derive CD200AR3&4KO (CD200R1&2 expressing) cells and (f) CD200AR4KO cells were used to make CD200 R1&4KO (CD200AR2&3expressing) cells. PCR validation for gene knockout for development of (hg) duel expressing and (h).

